# Gene function prediction in five model eukaryotes based on gene relative location through machine learning

**DOI:** 10.1101/2021.08.27.457944

**Authors:** Flavio Pazos Obregón, Diego Silvera, Pablo Soto, Patricio Yankilevich, Gustavo Guerberoff, Rafael Cantera

## Abstract

**Motiviation:** The function of most genes is unknown. The best results in gene function prediction are obtained with machine learning-based methods that combine multiple data sources, typically sequence derived features, protein structure and interaction data. Even though there is ample evidence showing that a gene’s function is not independent of its location, the few available examples of gene function prediction based on gene location relay on sequence identity between genes of different organisms and are thus subjected to the limitations of the relationship between sequence and function.

**Results:** Here we predict thousands of gene functions in five eukaryotes (*Saccharomyces cerevisiae, Caenorhabditis elegans, Drosophila melanogaster, Mus musculus* and *Homo sapiens*) using machine learning models trained with features derived from the location of genes in the genomes to which they belong. To the best of our knowledge this is the first work in which gene function prediction is successfully achieved in eukaryotic genomes using predictive features derived exclusively from the relative location of the genes.

**Supplementary information:** http://gfpml.bnd.edu.uy

## 1. INTRODUCTION

We witness a growing gap between the number of assembled genomes and the number of genes with known functions. Less than 1% of the protein sequences in UniProtKB (UniProt Consortium 2018) have an experimental Gene Ontology annotation (Ashburner et al. 2000) and even in well studied organisms, the majority of known genes have yet no assigned function (Zerbino et al. 2018). Furthermore, well studied genes have frequently been assigned more than one function so less studied genes, for which only one function is known, have probably more functions to be discovered (Rubin and Green 2013). It would take centuries to experimentally confirm the functions of the already known genes, hence the need to improve automatic functional prediction (AFP) (Bernardes and Pedreira 2013; Libbrecht and Noble 2015; Zhou et al. 2019; Zhao et al. 2020; Bonetta and Valentino 2020).

The Critical Assessment of protein Function Annotation algorithms (CAFA) is a series of experiments designed to provide a large-scale assessment of computational methods dedicated to automatic function prediction (AFP) (Radivojac et al. 2013; Jiang et al. 2016; Zhou et al. 2019). In all CAFA editions so far, the best results were obtained with machine learning-based methods combining multiple data sources, typically including sequence derived features, protein structure and molecular interaction data. The performance of the methods evaluated by the CAFA challenges improved dramatically between the first (2013) and the second (2016) edition but this improvement slowed down between the second and the third edition (2019). The authors hypothesized that including more varied sources of data will lead to additional large improvements in AFP (Zhou et al. 2019).

Thus, finding new ways to extract relevant biological information from the available data is key to improve AFP. For around 99% of all known proteins, the only available information is the sequence encoded in the corresponding genome, highlighting the importance of sequence-based AFP (Shehu et al. 2016). But AFP based on sequence similarity is hindered by a highly variable correlation between sequence identity and gene function (Duan et al. 2006) and by the evolutionary distance of many genomes to the closest well-characterized genome (Blaby-Haas and Merchant 2019). Here we explore the hypothesis that the relative location of a gene, a feature that is independent of sequence and can be directly extracted from any genome, is sufficient to perform AFP.

Functionally related genes may be constrained to remain close to each other due to natural selection, forming conserved gene clusters (Ling et al. 2009). Local clusters of co-expressed, co-regulated or functionally related genes have been documented in a wide range of organisms, including prokariotes, yeast, insects, vertebrates and plants (Eisen et al. 1998; Niehrs and Pollet 1999; Cohen et al. 2000; Boutanaev et al. 2002; Hurst et al. 2002; Lee and Sonnhammer 2003; Hurst et al. 2004; Michalak 2008).

Equating conserved co-locality with co-functionality have been a fruitful approach for predicting gene functions in prokaryotes for more than 20 years (Overbeek et al. 1999; Huynen et al. 2000; Wolf et al. 2001; Yanai et al. 2002; Zheng et al. 2002; Ling et al. 2009). On the contrary, there are very few examples (Mihelčić et al. 2019; Blaby-Haas and Merchant 2019) of the use of this approach in eukaryotic organisms, although also in eukaryotic genomes functions are non-randomly distributed (Lee and Sonnhammer 2003). However, these AFP studies were based on conserved gene neighborhoods, thus subjected to the limitations mentioned above regarding the relationship between sequence and function.

Here we performed AFP based exclusively on the relative location of genes. We test the predictive power of a feature which represents the spatial organization of genes with respect to their functions, which we term “functional landscape arrays” (FLAs). A FLA is an array associated to each gene that contains the enrichment in a set of Gene Ontology terms (GO terms) found around the gene, considering different window sizes. These arrays contain information which is independent of sequence similarity between genes and that can be automatically extracted from any annotated genome.

We predicted associations between genes of five eukaryotes (*Saccharomyces cerevisiae, Caenorhabditis elegans, Drosophila melanogaster, Mus musculus* and *Homo sapiens*) and terms from the three ontologies of the Gene Ontology (Biological Process, Cellular Component and Molecular Function) training a set of hierarchical multi-label classifiers with FLAs. Then we compared the results of our 15 models, one for each pair organism/ontology, with equivalent models that randomly assign functions to genes. We also compared -when possible-the performance of our models to the baseline methods used in CAFA 3: BLAST and Naïve Bayes. We found that our models, trained exclusively with location-derived features, performed several times better than chance and even better than BLAST when predicting Biological Process, showing that gene location can be more informative than sequence homology in order to obtain useful functional predictions.

## 2. SYSTEMS and METHODS

### 2.1 General procedure to predict associations between genes and GO terms

For each genome,

- Model as a string of protein coding genes.
- Random split in sets T and E, containing 80% and 20% of the genes respectively.

For each Ontology,

- Train a binary classifier for each GO term X associated with at least 40 genes in T and 10 genes in E
  - Training set: genes in T annotated with GO term X (as positives) and its siblings (as negatives)
  - Predictive feature: a FLA for each gene, including enrichment in GO term X, its siblings and its ancestors
  - Hyper-parameters set by grid search & cross validation
- Combine all the binary classifications into one hierarchical multi-label classifier using the node interaction method.
- Evaluate calculating the hF1 score over the test set E
- Using the classification threshold that maximizes hF1 over E, predict new associations between GO terms and genes in E.

### 2.2 Genome modeling

We model the genome as a collection of segments (the chromosomal arms) in which the protein coding genes -the only elements we considered-are located one next to the other, without intergenic regions or superpositions (Pazos Obregón et al. 2018). In this model, the position of a gene is defined by the location of its transcription starting point and the distance between two genes is the number of other genes located between them. The number of protein-coding genes considered in each genome is shown in Table 1.

**Table 1.** GO terms for which a binary classifier was trained and tested. The first column shows the assembly version used for each organism, the second column shows the number of protein coding genes in each genome, the third column indicates the ontology, the fourth column shows the number of GO terms associated with at least one gene for that organism and ontology and the fifth column shows the number of GO terms associated with at least 40 genes in the set T (used for training) and 10 genes in the set E (used for evaluation). These are the GO terms for which a binary classifier was trained and tested. For each organism and ontology, we implemented a hierarchical multilabel classifier combining these binary classifiers. The hierarchical precision, recall and F-max reached by each of these models are shown in columns sixth, seventh and eight.

| Organism | Protein coding genes | Ontology | Total GO terms | Considered GO terms | hPrec | hRec | hF-max |
| --- | --- | --- | --- | --- | --- | --- | --- |
| *S. cerevisiae*<br>(R64) | 5,892 | BP | 5,074 | 525 | 0.24 | 0.23 | 0.24 |
|  |  | CC | 1,035 | 137 | 0.51 | 0.52 | 0.52 |
|  |  | MF | 2,323 | 137 | 0.69 | 0.19 | 0.30 |
| *C. elegans*<br>(WBcel235) | 7,356 | BP | 5,661 | 551 | 0.09 | 0.15 | 0.11 |
|  |  | CC | 1,110 | 117 | 0.19 | 0.33 | 0.25 |
|  |  | MF | 2,226 | 151 | 0.25 | 0.14 | 0.17 |
| *D. melanogaster*<br>(BDGP6) | 11,122 | BP | 7,416 | 880 | 0.17 | 0.20 | 0.18 |
|  |  | CC | 1,277 | 176 | 0.41 | 0.37 | 0.39 |
|  |  | MF | 2,599 | 212 | 0.47 | 0.22 | 0.30 |
| *M. musculus*<br>(GRCm38.p6) | 20,809 | BP | 15,318 | 1040 | 0.22 | 0.21 | 0.21 |
|  |  | CC | 1,953 | 285 | 0.46 | 0.42 | 0.44 |
|  |  | MF | 4,269 | 364 | 0.63 | 0.25 | 0.36 |
| *H. sapiens*<br>(GRCh38.p13) | 17,276 | BP | 13,816 | 1212 | 0.21 | 0.20 | 0.20 |
|  |  | CC | 1,818 | 338 | 0.44 | 0.42 | 0.43 |
|  |  | MF | 4,244 | 369 | 0.47 | 0.27 | 0.35 |

### 2.3 Gene Ontology

Gene Ontology (GO) is an attempt to describe all the knowledge about the biological functions of genes with three ontologies: Molecular Function, Cellular Component and Biological Process, each one representing different aspects of the biology of a gene product and organized as a directed acyclic graph (Ashburner et al. 2000). Each “GO term” is a node of these graphs, with precise definition and relationships with other terms. A GO annotation occurs when an association between a gene product and a GO term is established. We used a version of the ontology downloaded on November 2018. To fulfill the true path rule (Valentini 2011), given the annotations of an organism within a given ontology, we up-propagated all the annotations, meaning that if a gene was annotated with a given GO term we associated that gene with all the ancestor terms up to the root of the graph.

### 2.4 Local enrichment analysis

Enrichment analysis is a method frequently used to determine if a given gene feature is overrepresented in a list of genes (Boyle et al. 2004). It assesses if the genes of a list associated with a given feature are more frequent than what should be expected in a list of genes of the same size but randomly picked from the same background list.

Given a gene of interest **j**, we define the Local Enrichment in the GO term **x** for the gene **j** and a window **w** centered in **j** as: **E**_**jxw**_ = ((**k**/**n**) / (**M**/**N**)), where **N** is the number of genes in the chromosomal arm, **M** is the number of genes in the chromosomal arm associated with GO term **x, n** is the number of genes in the window and **k** is the number of genes in the window associated with GO term **x** (see Figure 1). In other words, **E**_**jxw**_ assess if the genes annotated with the GO term **x** are located in the surroundings of gene **j** more frequently than what could be expected by chance. This approach was successfully used to look for clusters of GO terms along the genome of seven eukaryotes (Tiirikka et al. 2014).

**Figure 1.**
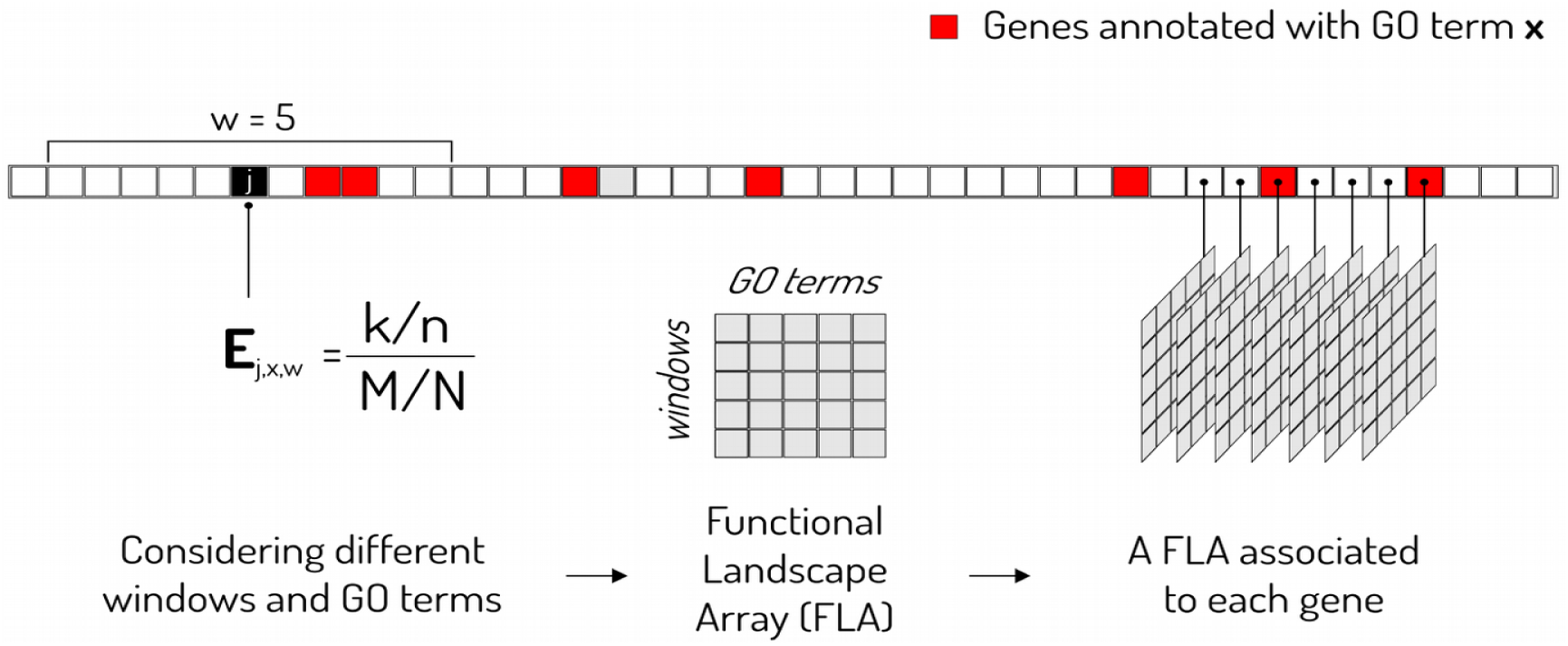
Local enrichment analysis and Functional Landscape Arrays. **k** is the number of genes in the window associated with GO term **x, n** is the number of genes in the window, **M** is the number of genes (squares) in the chromosomal arm (strip) associated with GO term **x**, and **N** is the total number of genes in the chromosomal arm.

### 2.5 Functional Landscape Arrays and Functional Enrichment Maps

To functionally characterize the surrounding of a gene we calculated its local enrichment in various GO terms. We considered a window **w**, centered in the gene under consideration, that includes 5, 10, 20, 50 or 100 genes to each side of the gene. The window was moved stepwise one gene at a time until the entire chromosome was covered (see Figure 1). Then, for each gene we defined a Functional Landscape Array (FLA): an array with a row for each window size and a column for each GO term whose enrichment was evaluated. Because of computational limitations, in the work we are reporting here, the GO terms included in each FLA depend on the GO term to be classified: we only included the enrichment found in that GO term, its father, its siblings and all its descendants.

Importantly: to train our models we did not consider the annotations of the genes in the set E, that was reserved for the evaluation of the models. This procedure guarantees an unbiased evaluation of the classifiers, in which the features used in training are not extracted from examples used in testing. Nevertheless, because it is a useful result by itself, we also performed Local Enrichment Analysis along each genome considering all its current annotations. We calculated the local enrichment around all the genes in each genome using the same set of window sizes and for all those GO terms associated with at least 20 genes and obtained what we call “functional enrichment maps”. The functional enrichment map of a given GO term shows which regions of a genome are enriched in that GO term, for various windows sizes.

### 2.6 Implementation of hierarchical multi label classifiers

We implemented a hierarchical multi label classifier for each pair organism / ontology using, with some modifications, the algorithm proposed in (Feng et al. 2017, 2018). This is a local approach, since a binary classifier is trained for each GO term. Due to computational limitations, for the binary classification at each node, instead of a Support Vector Machine, we used a Random Forest classifier (Breiman 2001), that have comparable performance in gene function prediction but with lower computational cost. For the same reason we did not used SMOTE (Chawla et al. 2002), a technique used to artificially generate new labeled data when training sets are too small-Depth, number of trees and measure of impurity for each classifier were set by grid search and 3-fold cross validation.

First we randomly split the genome into two sets: **T** and **E**. Set **T** includes 80% of the genes and was used to define the training sets and to obtain the FLAs. Set **E** includes the remaining 20% of the genes and was used to evaluate the models. We trained a binary classifier for each GO term that was associated with at least 40 genes in **T** and at least 10 genes in **E**. Table 2 shows the amount of GO terms meeting these conditions in each organism and ontology, i.e. the GO terms that could be predicted.

**Table 2.** Ratio between the hF-max reached by the trained model and the hF-max reached by the corresponding random model over the set E for each possible pair organism/ontology.

| Organism | BP | CC | MF | Mean |
| --- | --- | --- | --- | --- |
| *S. cerevisiae* | 1.74 | 2.02 | 1.34 | 1.70 |
| *C. elegans* | 1.76 | 2.08 | 1.15 | 1.66 |
| *D. melanogaster* | 2.14 | 2.22 | 1.83 | 2.06 |
| *M. musculus* | 2.78 | 2.87 | 2.33 | 2.66 |
| *H. sapiens* | 2.43 | 2.46 | 2.57 | 2.49 |
| Mean | 2.17 | 2.33 | 1.84 |  |

To define the training set for each classifier we applied the siblings policy (Silla and Freitas 2011). We included as positive cases those genes associated with the GO term under consideration and as negative cases those genes associated with the siblings or uncles terms of the GO term under consideration and not to that term. Importantly, to construct the FLA associated to each gene, to be used as predictive feature, we only considered the annotations of the genes that belonged to **T**.

With each trained classifier we classified the genes in **E** and then post-processed the predictions using the node interaction method (Feng et al. 2018), to respect the restrictions imposed by the hierarchy of the ontology. Finally, we evaluated the performance of each hierarchical multi-label classifier using the hierarchical version of the F1 score. All calculations were carried out using ClusterUY (site: https://cluster.uy).

### 2.7 Evaluation of the models

To evaluate the performance of each trained model we used the complete set of annotations of the genes in **E**, that were not used in training. As evaluation metric we used the hierarchical version of the F1 score (hF1) proposed in (Kiritchenko et al. 2006) and used in the CAFA competitions. If we denote the true and false positives as TP and FN and the true and false negatives as TN and FN, Precision (Pre) and Recall (Rec) are defined as:

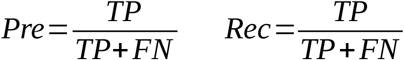

and their hierarchical versions, which we term hPre and hRec, are defined as:

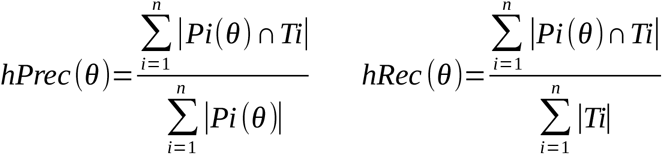

where θ ∈ [0,1] is the classification threshold, n is the number of genes, T*i* is the set of GO terms truly associated to gene *i* and P*i*(θ) is the set of GO terms predicted for gene *i* with the classification threshold set at θ. We assumed that the root of each ontology always is in P*i*(θ). The hF1 score is the harmonic mean of hPre and the hRec and is defined as:

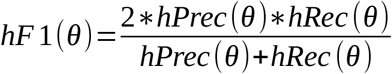

### 2.8 Comparison with random models

As a way to assess how far from randomness is the distribution of gene functions along the genome, we compared the hF1 of each of our trained models with the hF1 reached by an equivalent model that randomly assigns GO terms to genes. In these “random models”, the probabilities of association between each gene of the corresponding genome and each GO term for which a binary classifier was trained are randomly selected from a uniform distribution. For each organism and ontology, we obtained the ratio between the hF1 of the trained model and the hF1 of its random version.

### 2.9 Comparison to CAFA baseline methods

We also compared the performance of our models to that of the baseline methods used in CAFA 3: Naïve Bayes and BLAST. The Naïve method simply predicts the frequency of a term being annotated in a database (Clark and Radivojac 2011). BLAST was based on search results using the Basic Local Alignment Search Tool software against the training database (Altschul et al. 1990). A term was predicted as the highest local alignment sequence identity among all BLAST hits annotated with the term. Both of these methods were evaluated during CAFA 3, using the new experimental annotations accumulated during the competition, from February 2017 to November 2017. We used the same approach to compare the performance of these two CAFA methods to our method and used the new experimental annotations accumulated from November 2018 (date of the GO annotations that we had used to train our models) to September 2021.

We compared our models with the baseline methods when predicting GO terms for individual species. This data is available as Supplementary files for CAFA 3 at: https://doi.org/10.6084/m9.figshare.8135393.v3 and includes performance evaluation for *H. Sapiens, M. musculus* and *D. melanogaster*. We compared our results with those obtained with the limited-knowledge benchmarks and under the full evaluation mode. For more details about the different CAFA evaluations modes please refer to CAFA 3, Additional file 1 (Zhou et al. 2019) and CAFA2 (Jiang et al. 2016)

## 3. IMPLEMENTATION

### 3.1 Functional enrichment maps in 5 model eukaryotes

We performed Local Enrichment Analysis around each gene of a given genome considering windows of various sizes (See Methods). Local Enrichment Analysis of a given gene assess if the genes in the surroundings are annotated with any GO term more frequently than what could be expected by chance. Given a GO term, its functional enrichment map shows which regions of a genome are enriched in that GO term, considering various windows sizes. We obtained the functional enrichment map of all those GO terms associated with at least 20 genes in each of the five considered organisms. As an example, Figure 2 shows the functional enrichment maps in *D. melanogaster* of three GO terms that belong to the same branch of the Cellular Component ontology. The data to generate all the functional enrichment maps is available at: https://github.com/IIBCE-BND/gfpml-datasets/tree/master/lea

**Figure 2.**
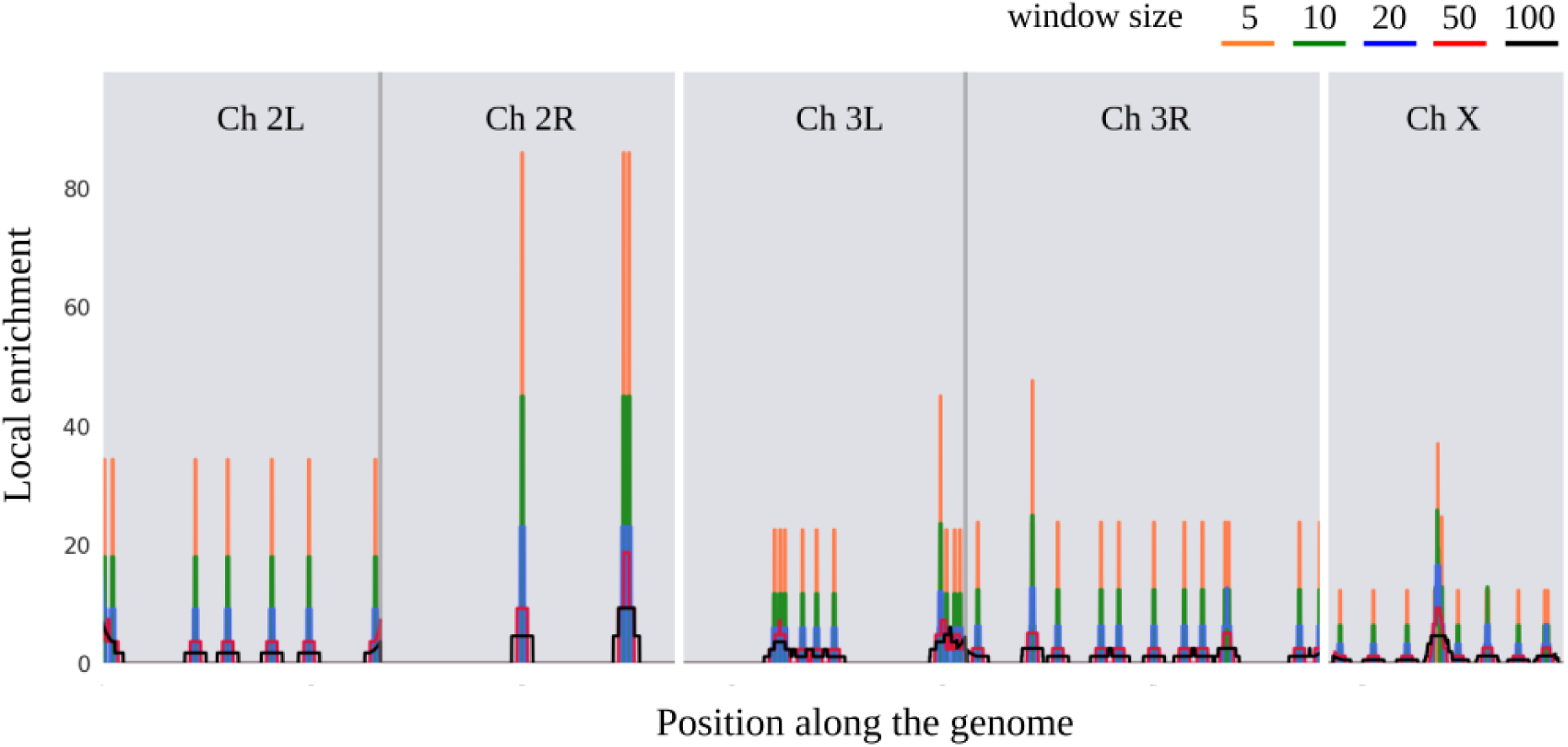
Functional enrichment map of the GO term “Golgi membrane” (GO:0000139) in the genome of D. melanogaster. There are 50 Drosophila genes annotated with this GO term that belongs to the Cellular Component ontology. The chromosomal position is represented in the x axis and the corresponding local enrichment at each position is shown in the y axis. Each light gray block corresponds to a chromosome (only chromosomes 2, 3 and X are shown) and the vertical dark gray lines mark the position of the centromeres, which divide the chromosome 2 into arms 2L and 2R and chromosome 3 into arms 3L and 3R. The enrichment found using different windows is shown with the colors indicated in the figure.

### 3.2 Implementation of hierarchical multi label classifiers

We trained fifteen hierarchical multi label classifiers, one for each possible pair organism/ontology. As detailed in Methods, we randomly split each genome into two sets: **T**, that includes 80% of the genes and was used for training, and **E**, that includes the remaining 20% of the genes and was used for evaluation. Each model assigned probabilities of association between the genes of the set **E** and those GO terms associated with at least 40 genes of the set **T** and 10 genes of the set **E**. Table 1 shows, for each organism and each ontology, the number of GO terms fulfilling these conditions and for which we implemented a binary classifier.

### 3.3 Evaluation of the models

We evaluated the performance of our models using the hierarchical version of the F1 score (hF1). Figure 3 shows the hF1 reached by each trained model over the test set **E**, as well as the hF1 of the corresponding random model, as a function of the classification threshold.

**Figure 3.**
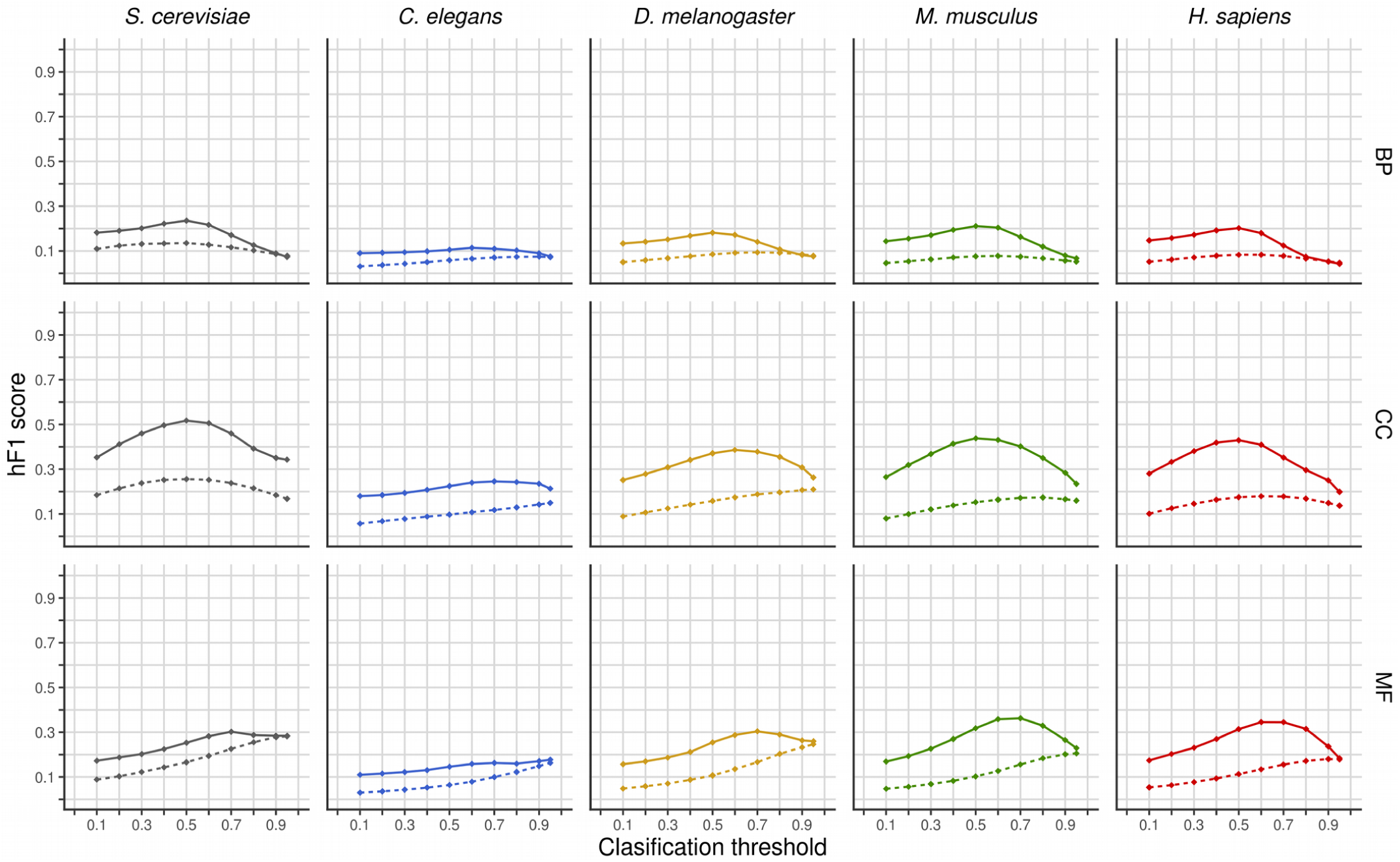
Hierarchical F1 over the test set for each model as a function of the classification threshold. In each plot the classification threshold, ranging from 0 to 1, is depicted in the x axis and the hF1, also ranging from 0 to 1, is depicted in the y axis. Trained models are represented by solid lines and random models by dotted lines. Each column of the panel corresponds to an organism and each row to an ontology (BP: Biological Process, CC: Cellular Component, MF: Molecular Function).

The hF-max is the highest hF1 score that the model reaches when varying the classification threshold. hF-max is a measure of the overall performance of the model and the corresponding classification threshold was used to predict new associations between GO terms and genes. Table 1 shows the hF-max for each model along with the corresponding precision and recall.

### 3.4 Comparison with random models

To assess how far from randomness is the linear organization of the genes along the genome with respect to its functions we calculated the ratio between the hF-max of the trained model and the hF-max of an equivalent random model, i.e. a model that randomly assigns probabilities of association between the same set of GO terms and the same genes (see Methods). Figures 4 and 5 show how this ratio varies with the classification threshold in each organism and ontology. Table 2 shows the hF-max reached by each model over the test set **E**. The trained models consistently performed better than the random models.

**Figure 4.**
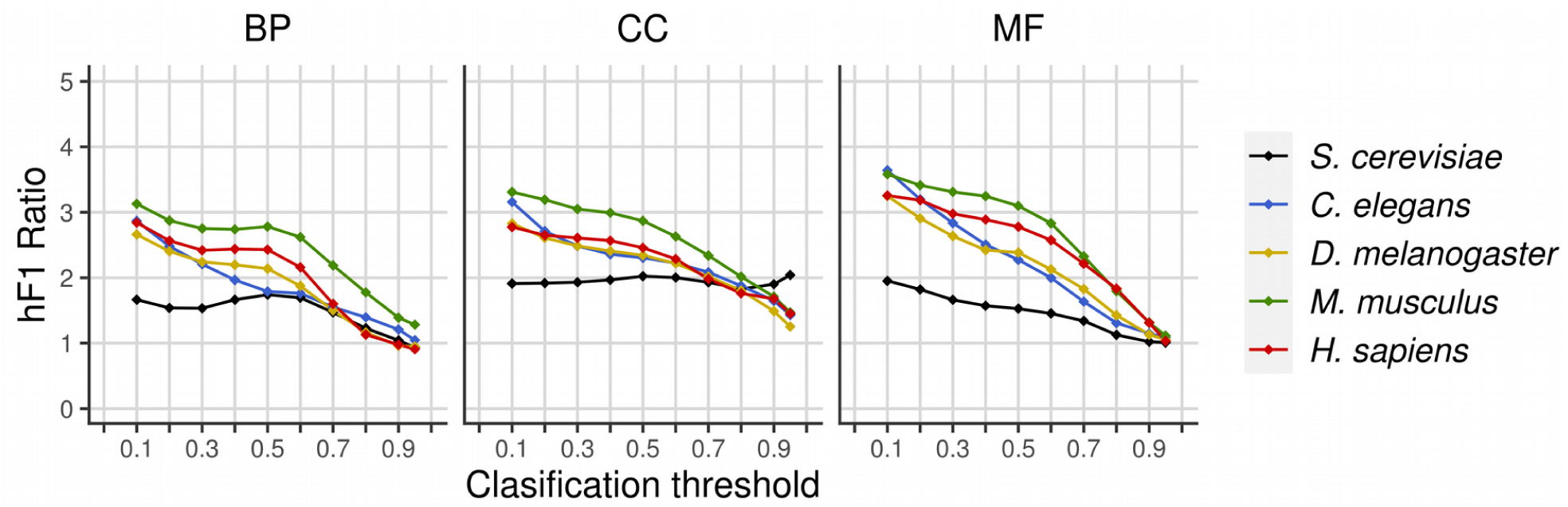
Ratio between the hF1 score of the trained model and the hF1 score of the corresponding random model as function of the classification threshold. Each graph shows the results for a given ontology, representing each organism with a different color.

**Figure 5.**
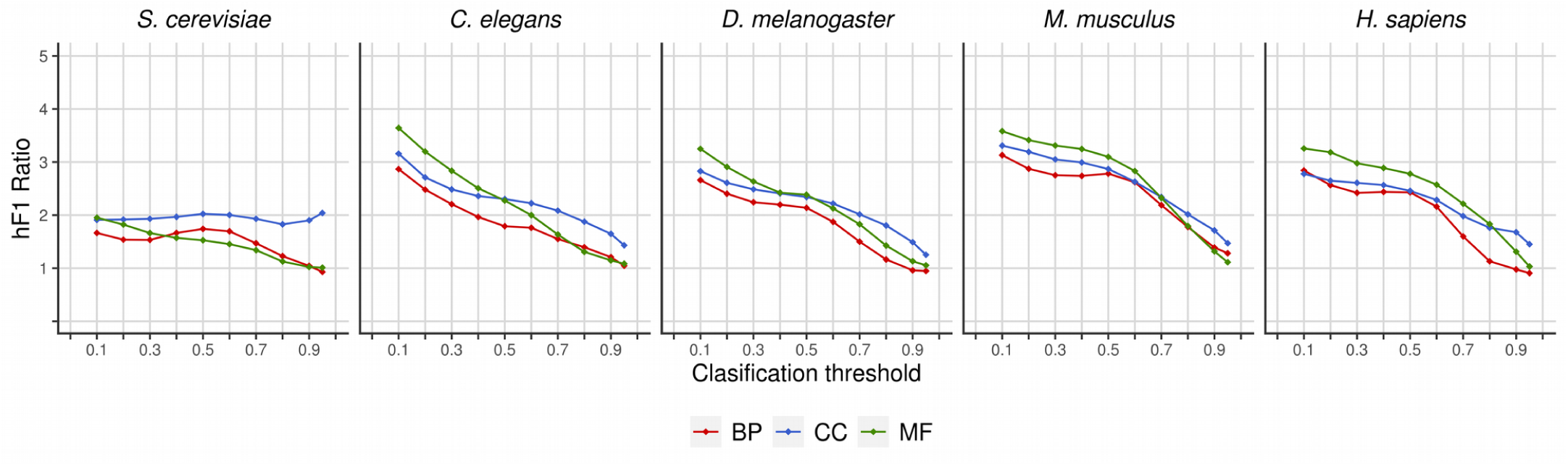
Ratio between the hF1 score of the trained model and the hF1 score of the corresponding random model as function of the classification threshold. Each graph shows the results for a given organism, representing each Ontology with a different color.

### 3.5 Comparison to CAFA baseline methods

As a complementary way to evaluate our models we also compared their performance with the performance reached by the baseline methods BLAST and Naïve Bayes used in CAFA 3 (see Methods). Comparisons were made between the hFmax reached by our models and the hFmax reached by BLAST and Naïve Bayes when making predictions for individual species, data that is only available for three of the five species we studied here: *H. sapiens, M. musculus* and *D. melanogaster*.

With this comparison we aim to asses if gene location alone can predict gene function with comparable performance to that reached by sequence homology or term frequencies alone. We found that this is the case and figure 6 shows the hFmax reached by the three models for each organism and ontology. Notably, for the three considered organisms the models trained with FLAs outperforms BLAST when predicting GO terms from the Biological Process Ontology.

**Figure 6.**
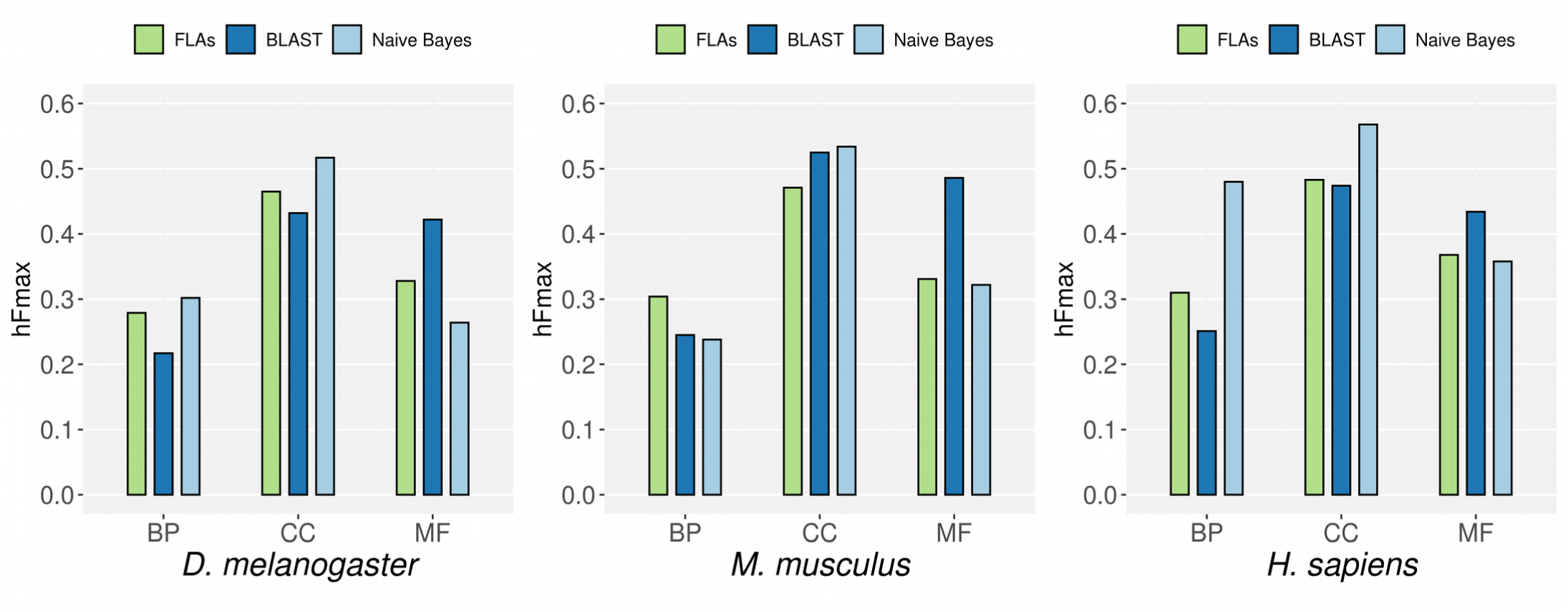
Comparison to CAFA baseline methods. Each graph shows the hFmax of different models when predicting GO terms of the three ontologies in three organisms. In green, the hFmax of the models exclusively trained with FLAs, evaluated using the new experimental annotations accumulated from November 2018 to September 2021. In blue and cyan, the hFmax of the baseline methods of CAFA 3 when making predictions on the same organisms and ontology as reported in (Zhou et al. 2019).

### 3.6 Prediction of new associations between genes and GO terms

We classified the genes in set **E** using the trained models and the classification threshold that maximizes the hF1 score (see Methods). We obtained the probability of association between each gene and each GO term associated with at least 40 genes in **T** and 10 genes in **E**. The complete set of predicted associations with a probability above the threshold is available at: http://gfpml.bnd.edu.uy. In this site, the user can browse and download all the predictions, searching by organism, ontology, chromosomal position, gene or GO term. Figure 7 shows a screenshot of the site.

**Figure 7.**
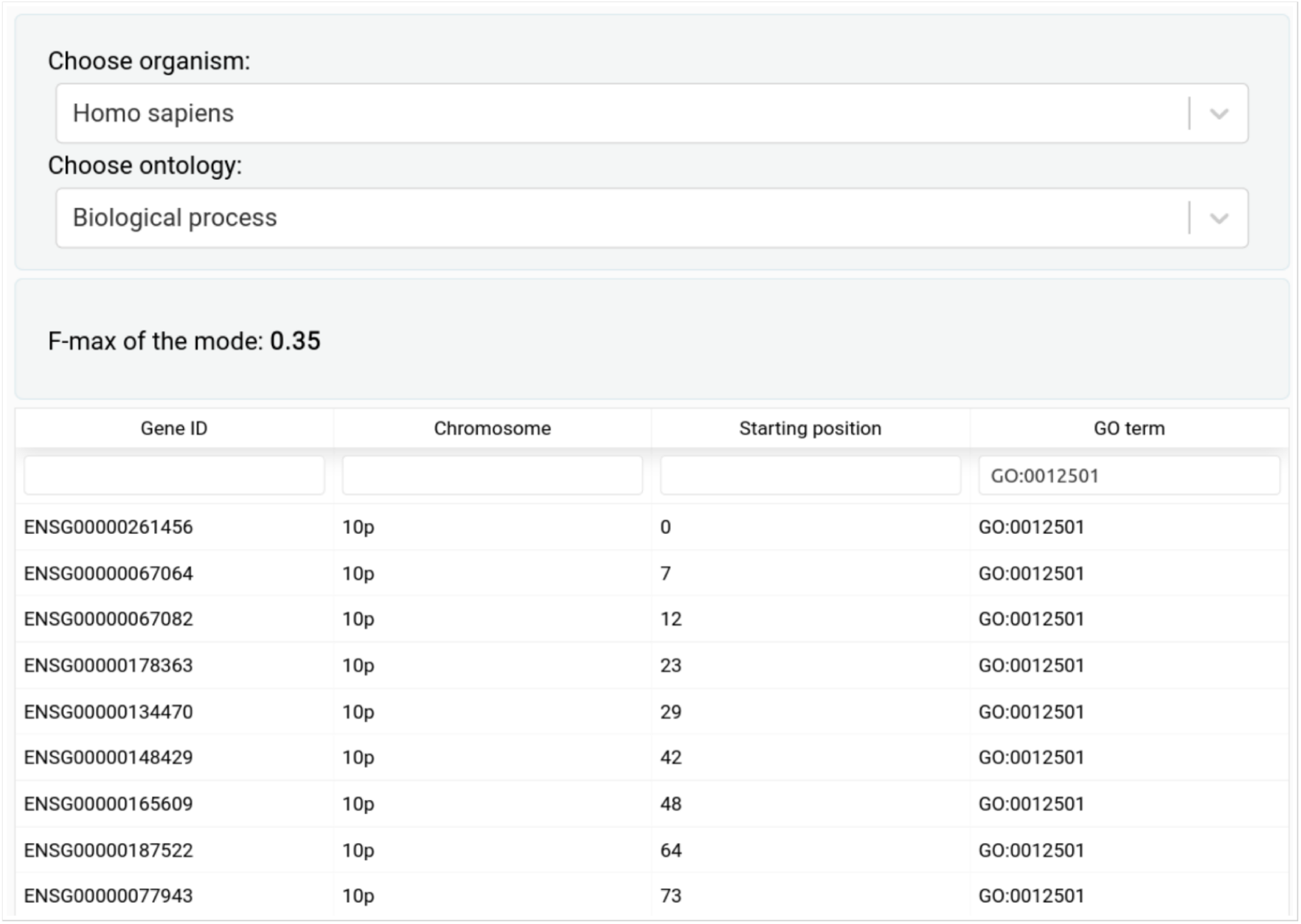
Screenshot of http://gfpml.bnd.edu.uy. At this website all the associations between genes and GO terms predicted by our models are available. Once an organism and an ontology is chosen, the hFmax reached by the corresponding model as well as the list of predicted associations are shown.

## 4. DISCUSSION

For the majority of the known genes, the only available information is their DNA sequence (Shehu et al. 2016). AFP based on DNA sequence similarity is a common approach, since its known that two genes with very similar sequences probably share function. But the contrary is not always true. A thorough study of the correlation between similarity in protein sequence and function in yeast (Duan et al. 2006) found that the majority of the sequences of proteins annotated with the same GO term were non-similar. In general, within one branch of an ontology tree, the more specific a GO term is, the more similar the sequences of the genes annotated with that term are, but the degree of similarity is highly variable and is significant only for specific GO terms. When using orthology between genes, these methods face another limitation: the evolutionary distance of many genomes to the closest well-characterized genome. For example, only 25–50% of the proteins in any given algal genome have detectable sequence similarity to any defined domain in the Pfam database (Blaby-Haas and Merchant 2019).

Spatial organization of genes (i.e. their localization along the genome) provides an alternative and complementary source of information that is independent of primary sequence (Ling et al. 2009). Genomic context-based methods, including gene neighborhoods, gene-order and gene-teams based methods, are a way to make use of this information (Shehu et al. 2016). These methods rely on orthology between genes and thus are subject to the above exposed limitations. Probably because these limitations, the few examples of genomic context-based AFP in eukaryotes are limited to a small proportion of the genes of the organism being considered (Mihelčić et al. 2019; Foflonker and Blaby-Haas 2020).

There is plenty of evidence pointing to the existence of distinctive patterns in the way in which functionally related genes distribute along eukaryotic genomes. If such patterns exist and are biologically relevant, it should be possible, at least in some cases, to predict the functions of a gene using as predictive feature its relative position with respect to other genes of known function in the same genome. As far as we know, here we have performed this task for the first time, using a new way to represent the information contained in these patterns: the Functional Landscape Arrays. This feature can be automatically extracted from any annotated genome and does not depend on orthology relations with other organisms.

Using FLAs as the only predictive feature we trained a set of hierarchical multi label classifiers that outperforms BLAST when predicting GO terms from the Biological Process Ontology in *H. sapiens, M. musculus* and *D. melanogaster* (see Figure 6). Our models also outperform BLAST when predicting GO terms from the Cellular Component Ontology in *H. sapiens* and *D. melanogaster*. With regard to the Molecular Function Ontology, our models have a performance that is similar to that of Naïve Bayes. With these trained classifiers we obtained thousands of new predicted associations between genes and GO terms in five eukaryotes, available at *http://gfpml.bnd.edu.uy*.

The relevance of our results stems from the fact that the performance of our models, assessed by standard metrics, shows that AFP exclusively based on features derived from the relative location of genes can be successfully performed on eukaryotic genomes. Even though in AFP is usual to integrate multiple types of information, information derived from gene location is rarely taken into account. Furthermore, according to the CAFA organizers, new improvements in gene function prediction should be expected from the incorporation of new kinds of predictive features (Zhou et al. 2019). We believe that including FLAs as predictive feature could significantly improve the performance of AFP models.

Our results are interesting from another point of view. The existence in eukaryotes of distribution patterns of functionally related genes so well defined as to allow good AFP points to levels of organization thought to be exclusive of prokaryotic genomes and its characteristic operons (Diament and Tuller 2016) Diament and Tuller performed a comparative study of the organization of several genomes, analyzing the location of functionally related genes. Their results revealed that the prokaryote *Escherichia coli* exhibits a higher level of genomic organization than the eukaryote *S. cerevisiae*, as one would expect given its operon-based genomic organization. But when considering a higher order of genomic organization, analyzing the co-localization of pairs of different functional gene groups, the authors found that the genome of *S. cerevisiae* is markedly more organized than that of *E. coli*. Our results are consistent with this trend. To estimate how far from randomness is the linear organization of different genomes we used the hF-max ratio, i.e. the ratio between the hF-max reached by the trained model and the hF-max reached by the random model. Table 2 and Figure 4 show that although the relationship between the complexity of the organism and its hF-max ratio is not linear, simpler organisms reach lower hF-max ratios than more complex organisms.

In sum, Functional Landscape Arrays have the potential to improve AFP, as they can be easily integrated into any model, can be automatically extracted from any annotated genome and are independent from sequence identity. To the best of our knowledge this is the first work in which only features derived from the relative gene location of the genes within a genome are used to successfully predict gene function in eukaryotes.

## Funding

This work was supported by Agencia Nacional de Investigación e Innovación, Uruguay, [grant number FSDA_1_2017_1_14242]; Instituto de Investigaciones Biológicas “Clemente Estable”, MEC, Uruguay and Programa de Desarrollo de las Ciencias Básicas, Uruguay.

The experiments presented in this paper were carried out using ClusterUY (site: https://cluster.uy).

